# PepN is a non-essential, cell wall-localized protein that contributes to neutrophil elastase-mediated killing of *Streptococcus pneumoniae*

**DOI:** 10.1101/313569

**Authors:** Charmaine N. Nganje, Scott A. Haynes, Christine M. Qabar, Rachel C. Lent, Elsa N. Bou Ghanem, Mara G. Shainheit

**Affiliations:** Department of Biological Sciences, Towson University, Towson, MD, USA; Department of Microbiology and Immunology, University at Buffalo School of Medicine, Buffalo, NY, USA

## Abstract

*Streptococcus pneumoniae* (*Spn*) is an asymptomatic colonizer of the human nasopharynx but can also cause invasive diseases in the inner ear, meninges, lung and blood. Although various mechanisms contribute to the effective clearance of *Spn*, opsonophagocytosis by neutrophils is perhaps most critical. Upon phagocytosis, *Spn* is exposed to various degradative molecules, including a family of neutrophil serine proteases (NSPs) that are stored within intracellular granules. Despite the critical importance of NSPs in killing *Spn*, the bacterial proteins that are degraded by NSPs leading to *Spn* death are still unknown. In this report, we identify a 90kDa protein in a purified cell wall (CW) preparation, aminopeptidase N (PepN) that is degraded by the NSP, neutrophil elastase (NE). Since PepN lacked a canonical signal sequence or LPxTG motif, we created a mutant expressing a FLAG tagged version of the protein and confirmed its localization to the CW compartment. We determined that not only is PepN a *bona fide* CW protein, but also is a substrate of NE in the context of intact *Spn* cells. Furthermore, in comparison to wild-type TIGR4 *Spn*, a mutant strain lacking PepN demonstrated a significant hyper-resistance phenotype *in vitro* in the presence of purified NE as well as in opsonophagocytic assays with purified human neutrophils *ex vivo*. Taken together, this is the first study to demonstrate that PepN is a CW-localized protein and a substrate of NE that contributes to the effective killing of *Spn* by NSPs and human neutrophils.

**IMPORTANCE:** Neutrophils are innate immune cells needed to effectively clear *Streptococcus pneumoniae* (*Spn*). Neutrophil serine proteases (NSPs) are important for killing phagocytosed *Spn*, however, the identity of the *Spn* proteins that are degraded by NSPs are unknown. This study identifies a *Spn* cell wall protein, aminopeptidase N (PepN) that is degraded by the NSP, neutrophil elastase (NE). We demonstrate that PepN is a *bona fide* cell wall protein and mutants lacking PepN are significantly more resistant than wild-type to killing by purified NE and human neutrophils. This study demonstrates that PepN is a NE substrate and its degradation contributes to effective *Spn* killing. By better understanding how neutrophils kill *Spn*, we aim to inform the development of improved therapeutic interventions.

## INTRODUCTION

*Streptococcus pneumoniae* (*Spn*) is a Gram-positive bacterium that is a frequent, asymptomatic colonizer of the human upper respiratory tract. However, if it gains access to sterile sites in the human host, such as the lungs, inner ear, meninges or blood, it can cause a variety of invasive diseases including pneumonia, otitis media, meningitis and sepsis, respectively [1-4]. Due to these invasive infections, about 1 million children per year under the age of five die, mostly in the developing world where access to healthcare is limited [5]. Neutrophils are the most abundant white blood cell in the body and are often the first immune cell type to migrate to the site of infection [6, 7]. Neutrophils play a critical role in the effective clearance of *Spn* via the process of opsonophagocytic killing. This multi-step process involves the tagging of *Spn* cells with complement proteins and subsequent internalization and degradation through the action of various factors including reactive oxygen and nitrogen species, antimicrobial peptides and a family of enzymes contained within the azurophilic granules, neutrophil serine proteases (NSPs) [8]. Of this repertoire of anti-microbial factors, previous work demonstrated that NSPs are the most important component for effectively killing *Spn in vitro* [9] and play a vital, protective role in murine models of pneumococcal pneumonia [10]. Furthermore, in individuals with Chediak-Higashi syndrome, a rare genetic disorder that impairs the mobilization of NSP-containing granules [11], neutrophils exhibited a reduced ability to kill *Spn* [12].

To date, four enzymes have been identified as members of the NSP family, neutrophil elastase (NE), cathepsin G (CG), proteinase 3 (PR3) and neutrophil serine protease 4 (NSP4) [13, 14]. NSPs are members of the chymotrypsin family of serine proteases and contain a His-Asp-Ser catalytic triad [14, 15]. NSPs become enzymatically active when granule-containing NSPs fuse with the phagocytic compartment and can also be exocytosed as a component of neutrophil extracellular traps (NETs) to combat extracellular pathogens [16, 17]. Several studies revealed that NSPs can reduce bacterial pathogenicity by destroying virulence factors produced by a range of pathogens, including: *Shigella flexneri, Salmonella enterica* serovar Typhimurium*, Yersinia enterocolitica* and *Staphylococcus aureus* [18, 19]. In addition, NSPs have been shown to directly kill *Pseudomonas aeruginosa*, *Escherichia coli* and *Klebsiella pneumoniae* [20-23]. Specifically, in *E. coli*, it was revealed that NE degrades OmpA, which destabilized the cell and induced cell death [20, 21]. Additionally, NE-mediated degradation of OprF, a major outer membrane protein in *P. aeruginosa*, was demonstrated to be necessary for effective immune defense in a mouse model of lung infection [22]. These findings emphasize the importance of NSPs in anti-microbial defenses and that they achieve this, in part, by degrading specific bacterial proteins. Despite the importance of NSPs in controlling *Spn* infection [9, 10], the exact surface proteins on this pathogen that are degraded by NSPs leading to *Spn* death have yet to be identified.

In this study, we aimed to identify specific cell wall (CW)-localized *Spn* proteins that are degraded by NE and/or CG, since these two NSPs were shown to be important for killing *Spn* both *in vitro* and *in vivo* [9, 10]. In experiments using a purified CW preparation, we identified a ~90kDa protein that was specifically and significantly degraded by purified NE. Analysis by mass spectrometry revealed this protein to be aminopeptidase N (PepN), an annotated metalloproteinase predicted to cleave a variety of peptides from the N-terminus [24]. Since PepN lacked an obvious secretion signal or CW localization motif, we epitope-tagged the C-terminus and performed sub-cellular fractionation experiments that revealed that PepN did indeed localize to the CW compartment. Importantly, PepN was shown to be a substrate of purified NE both in experiments with purified CW and in intact *Spn* cells. Furthermore, a mutant strain of *Spn* lacking PepN (Δ*pepN*) was significantly more resistant to killing by purified NE and opsonophagocytic killing by human neutrophils. Taken together, these data identify the first *Spn* cell wall protein that is degraded by the NSP, neutrophil elastase, and demonstrate that the degradation of PepN contributes to the effective killing of *Spn*.

## MATERIALS AND METHODS

### Bacterial strains, growth conditions and growth curves

*S. pneumoniae* strain TIGR4 (serotype 4) was used throughout this study. *S. pneumoniae* was grown in Todd-Hewitt broth (BD) supplemented with 0.5% yeast extract (Fisher) (THY) and oxyrase (5μL/mL) or trypticase soy broth (TSB) (BD) supplemented with catalase (Sigma; 30U/mL). Alternatively, bacteria were grown on tryptic soy agar (TSA) supplemented with 200U/mL of catalase. Where appropriate during growth, antibiotics at the following concentrations were included: chloramphenicol (Cm; 4μg/mL) or streptomycin (Sm; 100μg/mL). All cells were grown at 37°C in a 5% CO_2_ incubator.

For growth curve experiments, strains of interest were grown to mid-log phase (OD_600_ ~0.6) in THY supplemented with oxyrase and then back-diluted to an OD_600_ ~0.003 in fresh media. Next, 200μL of cells were added to 96 well flat bottom plates in replicates of 8 wells per strain and were incubated at 37°C for 12h. OD_600_ measurements were taken every 30 minutes using a VersaMax plate reader.

### Generation of mutant strains

All mutant strains of *S. pneumoniae* used in this study are described in **Table 1** and were generated by allelic exchange using the DNA constructs shown in **Figure 1A.** Each allelic exchange construct was generated *in vitro* using splicing by overlap extension PCR [25]. The upstream and downstream arms of homology flanking *pepN* were PCR amplified from TIGR4 genomic DNA. For Cm^r^ mutant strains, the chloramphenicol resistant cassette was amplified from pAC100 [26]. To make the PepNFLAG tag and the Δ*pepN*Revertant strains, we performed co-transformation with the mutant *rpsL* allele (Sm^r^) and the corresponding DNA construct to introduce a FLAG epitope directly upstream of the stop codon or restore the *pepN* gene **(Figure 1A)** [27, 28]. Transformation of *S. pneumoniae* was performed as previously described [29]. All mutations were confirmed by PCR and Sanger sequencing (Eton Biosciences).

**Table 1.**
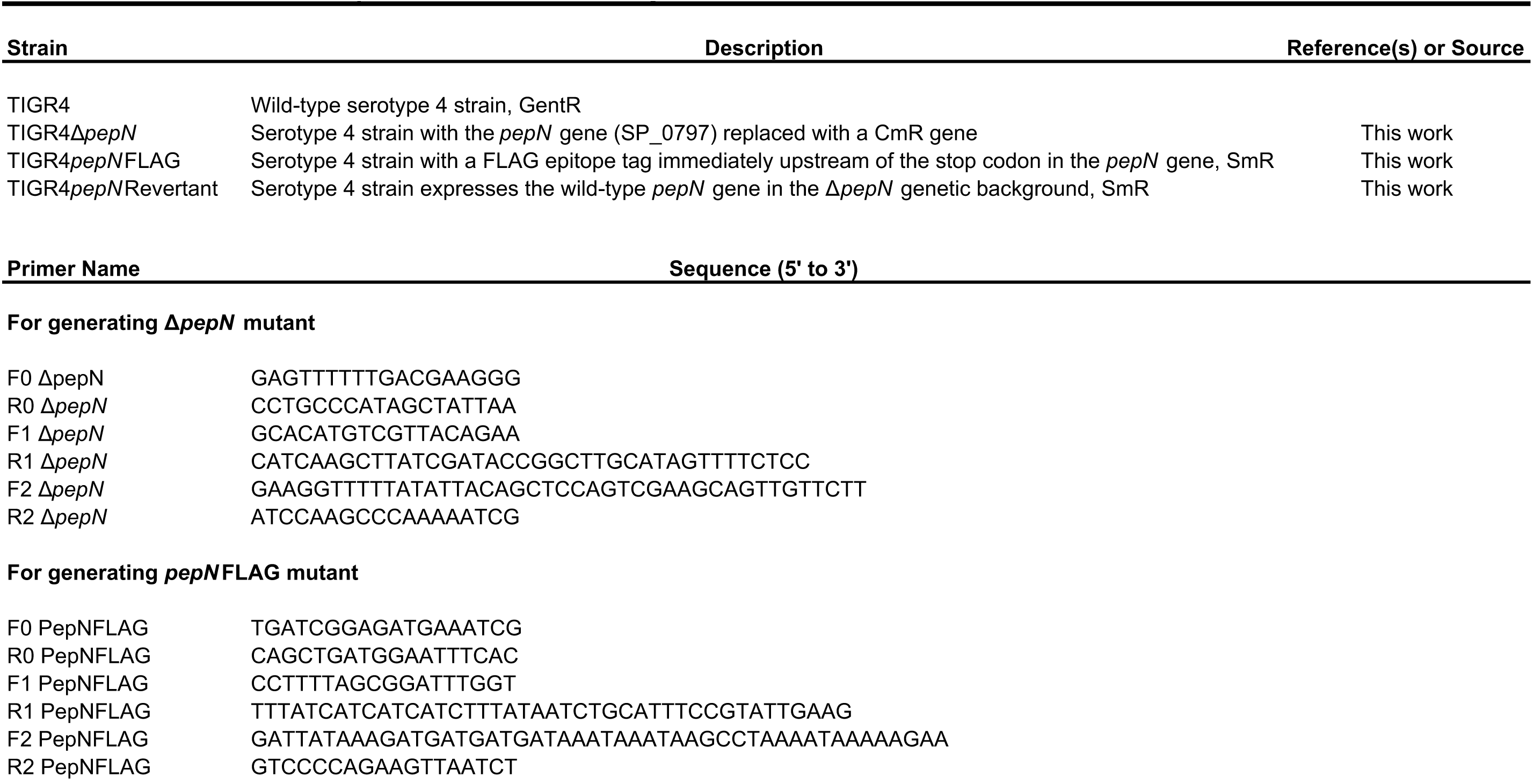
Relevant strains and primers used in this study

**Figure 1.**
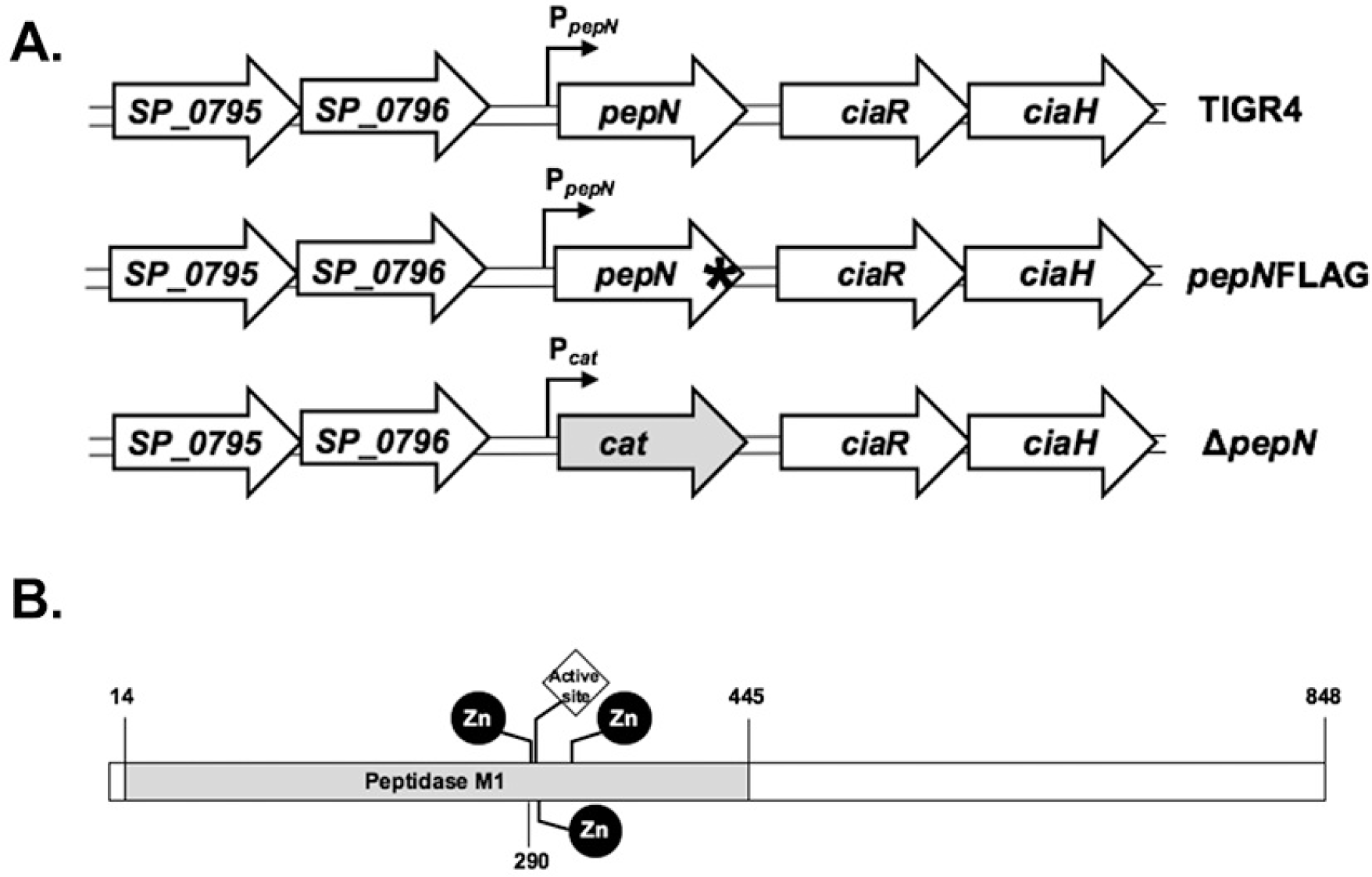
Schematic representation of the DNA constructs used create *Spn* mutant strains and of the domain structure of the PepN protein. (**A)** The depicted DNA constructs were used to create mutant strains of *S. pneumoniae* via allelic exchange. In the *ΔpepN* strain, the *pepN* ORF was replaced by a Cm resistance cassette. In the *pepNFLAG* mutant a 1X FLAG tag (denoted by *) was added to the C terminus of the PepN protein, immediately upstream of the stop codon. **(B)** The domain structure of PepN was determined using BLASTP to be comprised of a peptidase M1 domain containing an active site coordinated by three zinc binding sites (residues 292, 296 and 315).

### Sub-cellular fractionation of *S. pneumoniae*

To isolate purified cell walls and protoplasts, we followed an established protocol as described previously [30], with some modifications. Briefly, 10mL of mid-log cultures (OD_600_ ~0.6) were pelleted, washed once with 1mL 50 mM Tris-Cl, pH 7.5 and resuspended in 100 μL of cell wall digestion buffer (CWDB) containing 50 mM Tris-Cl, pH 7.5, 30% (w/v) sucrose, 1 mg/mL lysozyme, 300 U/μL mutanolysin (both Sigma-Aldrich), 1x protease inhibitor cocktail (Roche). Cells were incubated for at least 2 h at 37°C with rotation. Whole cell lysates (WCL) were prepared by directly adding SDS buffer to samples and boiling them for 10 mins at 100°C. For protoplast and cell wall (CW), samples were centrifuged at 13,000 × g for 10 min. The supernatant, containing the CW, was applied to SpinX 0.22μm spin columns (Sigma) and centrifuged for 1 min at 13,000 × g to remove any contaminating protoplasts. The pellet, containing the protoplasts, was resuspended in 100μL 50mM Tris-Cl pH 7.5. Samples were stored at −20°C until further use.

### Capsule immunodot blot assays

To quantify the amount of capsule present in wild-type TIGR4 and *ΔpepN* mutant strains, 1 ml of OD_600_-matched mid-exponential-growth-phase bacteria was pelleted and stored at −20°C until use. Samples were resuspended in 300 μl of CWDB, described above, and incubated at 37°C for 30 min. Samples were then sonicated for 4 – 10 second intervals while on ice using a probe sonic dismembrator (Fischer Scientific). Next, a 2-fold serial dilution of lysate was prepared in 1X PBS, and 5 μl was spotted on 0.2-μm-pore-size nitrocellulose membranes (Invitrogen, Inc.) with suction. Membranes were blocked with 10mL of 5% milk in 1X TBS with 0.1% Tween-20 (TBST) for 1 h at room temperature with shaking, then washed 1X with 10mL 1X TBST for 5 min. The membrane was then probed with an unconjugated rabbit anti-serotype 4 serum (Statens Serum Institut; 1:1000) in 5mL of 2.5% milk in 1X TBST for 1h at room temperature with shaking. After 3-5 min washes with 15mL 1X TBST, an HRP-conjugated goat-anti-rabbit secondary antibody (1:2500, Jackson ImmunoResearch) was applied to the membrane in 5mL of 2.5% milk in 1X TBST for 1h at room temperature with shaking. Finally, the membrane was washed 3X with 10mL 1X TBST, developed using ECL Blotting Substrate (Thermo Scientific), visualized using the C-Digit Western Blot Scanner, and quantified using ImageStudioLite (LI-COR Biosciences).

### Neutrophil elastase-cell wall degradation assays and SDS-PAGE analysis

CW purified from 10^8^ CFU were incubated with 68μM of neutrophil elastase (NE) (Elastin Products Company) or left untreated for 24 h at 37°C with rotation. Equal volumes of samples were boiled for 10 min in SDS sample buffer containing 50mM Tris-HCl, 10% glycerol, 2% SDS, 0.1% bromophenol blue, 2% β-mercaptoethanol and run on an AnykD gradient Tris-glycine polyacrylamide gel (Bio-Rad). Gels were stained with Colloidal Coomassie Blue (Bio-Rad) for 18h and were visualized using an Epson Perfection V550 Photo scanner. To determine the identity of proteins degraded by NE, bands of interest were excised from the gel and analyzed by the Taplin Mass Spectrometry Facility at Harvard University.

### Subcellular localization of PepN via Western blot analysis

CW was isolated from 10^7^ CFU of the appropriate bacterial strains. Equal volumes of samples were analyzed via SDS-PAGE as described above, proteins were transferred to a nitrocellulose membrane (ThermoFisher) at 22V for 18h at 4°C and blocked for 1 h at room temperature with 5% skim milk in 1X PBS with 0.2% Tween-20 (PBST). Unconjugated primary mouse anti-FLAG antibody (Sigma-Aldrich) at 1:3,000 and rabbit anti-CodY serum (a generous gift from Dr. A. L. Sonenshein) at 1:10,000 diluted in 2.5% milk in 1X PBST were added to membranes and incubated for 1 h at room temperature with agitation. Membranes were then washed 3X with 10mL 1X PBST for 5 mins each. HRP-conjugated secondary antibodies (Jackson ImmunoResearch) were applied at a 1:10,000 dilution in 2.5% milk in 1X PBST and incubated for 1 h at room temperature, followed by three 45 min washes in 10mL 1X PBST. Membranes were then developed with SuperSignal West Dura Extended Duration Substrate (ThermoFisher).

### *In vitro* bactericidal and PepN degradation assays with neutrophil elastase

Wild-type and mutant strains of *S. pneumoniae* were grown to early-exponential phase (OD_600_ ~0.3) in TSB supplemented with catalase, pelleted by centrifugation, washed in sterile 1X PBS, and resuspended in 10 μM sterile sodium phosphate reaction buffer to a concentration of ~10^7^ CFU/mL. 50μL of cells were exposed to a two-fold dilution series of NE ranging from 3.4 – 13.6μM and incubated for 1h at 37°C with 5% CO2. To determine viable counts, cells were then serially diluted, plated on TSA and incubated overnight at 37°C with 5% CO_2_.

To assess whether PepN is degraded by NE in the context of an intact *Spn* cell, ~5 × 106 CFU were exposed to 6.8μM NE, or left untreated, for 1h at 37°C with 5% CO_2_, followed by the isolation of CW and WCL fractions as described above. Samples were then subjected to SDS-PAGE and Western Blot analysis using anti-FLAG or anti-CodY antibodies. Data were compiled from at least three independent experiments.

### *Ex vivo* neutrophil opsonophagocytic killing assay

Young, healthy human volunteers were recruited in accordance to Tufts Medical Center Human Investigation Review Board (IRB) and signed informed consent forms. Individuals taking medication, reporting symptoms of infection within the last 2 weeks or that were pregnant were excluded from the study. Whole blood was obtained and anticoagulated with acid citrate/dextrose. PMNs were isolated using a 2% gelatin sedimentation technique as previously described [31] which allows for isolation of active PMNs with ~90% purity. Killing of opsonized *Spn* by human neutrophils was performed as previously described [32] based on a modified protocol described by Dalia *et al.* [33]. Strains of *Spn* were grown to mid-log and 10^3^ CFU were opsonized in 10% (v/v) baby rabbit serum (Pel-Freeze) in 200μL of Hank’s Buffer supplemented with 0.1% gelatin for 30 mins. Pre-opsonized bacteria were then mixed with 5 × 10^5^ PMNs for 45 mins at 37°C with rotation. Samples were then placed on ice to stop the process of opsonophagocytosis followed by serial dilution and plating to enumerate viable CFU. The percentage of bacterial killing was calculated relative to controls without PMNs.

### Statistical analysis

To determine if a statistically significant difference existed between wild-type TIGR4 and the mutant strain of interest across various concentrations of NE, two-way ANOVA statistical tests were performed [34]. In experiments that compared the phenotypes of wild-type TIGR4, *ΔpepN* and *ΔpepNRev* strains in the presence of only 3.4 μM NE, a one-way ANOVA followed by a multiple comparisons test was performed. Unpaired Student t-test was used for comparison of bacterial killing by PMNs. *p* values less than 0.05 were considered significant. All statistical analyses were performed using GraphPad Prism for Mac (GraphPad Software, Inc).

## RESULTS

### Analysis of purified cell wall (CW) from TIGR4 treated with the NSPs, neutrophil elastase (NE) or cathepsin G (CG)

Although NSPs were demonstrated to be essential for the effective killing of *Spn* [9], the protein targets that are degraded by these enzymes remains unknown. To identify *Spn* CW proteins that are degraded by NSPs, CW was isolated from 10^8^ mid-log bacteria and were incubated with 68μM NE, 10μM CG or left untreated. Samples were then evaluated using SDS-PAGE analysis followed by staining with Coomassie blue. As depicted in **Figure 2A,** several bands **(arrow heads)**, most notably a 90kDa protein **(black arrows)**, were absent only in the sample treated with NE, but was clearly visible in both the untreated and CG-treated samples. In order to determine the identity of this specific NE substrate, we excised the 90kDa band from the gel in both the untreated and NE-treated samples for mass spectrometry analysis. This analysis revealed that the 90kDa band was vastly comprised of a single protein species (>90%), which is reported as the “percent intensity” value in **Figure 2B.** Importantly, based on its amino acid sequence, this protein was determined to be aminopeptidase N (PepN; SP_0797). Additionally, by comparing the relative abundance of the PepN protein in the untreated versus NE-treated samples, presented in **Figure 2B** as “Sum intensity”, we observed that PepN was substantially degraded, with up to a 10^4^-fold reduction in protein abundance. Furthermore, CW isolated from a mutant strain of *Spn* that lacks PepN (Δ*pepN*) did not possess the 90kDa band, confirming that the vast majority of that band was comprised of PepN **(Figure 2A)**. Taken together, these data demonstrate that PepN is a 90kDa protein found in a purified *Spn* CW preparation that is specifically and markedly degraded by NE.

**Figure 2.**
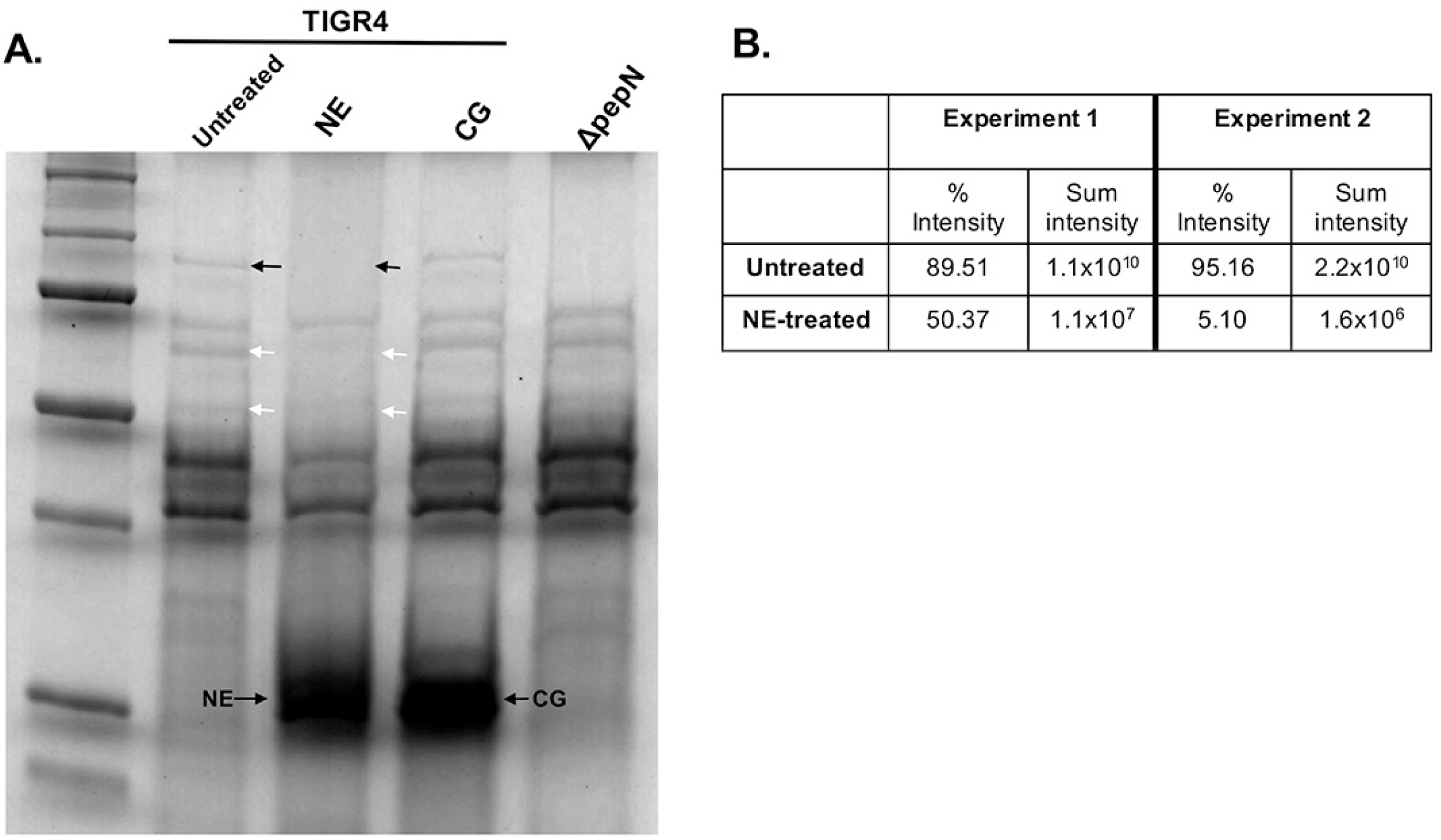
Neutrophil elastase degrades a 90kDa *Spn* cell wall (CW) protein *in vitro*. Neutrophil elastase degrades a 90kDa *Spn* protein in a purified cell wall fraction TIGR4 and Δ*pepN* strains of *S. pneumoniae* were grown to mid-log, 10^8^ cells were pelleted by centrifugation, washed in 1 X PBS and incubated with CWDB for 18h at 37°C with rotation, followed by centrifugation using a 0.22μm spin column to separate the CW and protoplast fractions. Purified CW samples were left untreated or incubated with 68μM NE or 10μM CG for 24h at 37°C with rotation. **(A)** Samples were then boiled in SDS sample buffer and analyzed via using an AnykD gradient Tris-glycine polyacrylamide gel followed by Coomassie blue staining. Arrow heads indicate bands degraded by NE. Black arrow heads highlight the 90kDa species degraded by NE. **(B)** To determine the identity of the 90kDa protein that is degraded by NE, we excised this band from both the untreated and NE-treated lanes and had it analyzed via mass spectrometry. Additionally, this analysis quantified the relative abundance of the 90kDa protein in both the untreated and NE-treated samples. The gel is from one experiment representative of 5 independent experiments. The mass spectrometry analysis is from 2 of those independent experiments.

### Characterization of PepN as a *bona fide* CW protein in *Spn* that is degraded by NE *in vitro*

Based on its amino acid sequence, PepN is annotated as a member of the peptidase M1 family and possesses an active site that is coordinated by three zinc binding sites **(Figure 1B)**. However, examination of the PepN amino acid sequence failed to reveal a canonical LPxTG motif, choline binding domain, Sec secretion signal peptide, or any other characterized export signal [35]. While it is not impossible for a protein lacking a classic export sequence to be secreted from the cell, such as the Pht proteins and those described as non-classical surface proteins [35, 36], this prompted a more thorough investigation of the sub-cellular localization of PepN. To do so, we generated a mutant strain of *Spn* via natural transformation using a DNA construct that added a FLAG epitope immediately upstream of the stop codon **(Figure 1A; *pepNFLAG*)**. For these experiments, we prepared CW, protoplasts and whole cell lysates (WCL) from mid-exponential phase TIGR4 and *pepNFLAG* cells and analyzed these fractions via Western Blotting using anti-FLAG or anti-CodY (a cytoplasmic protein) antibodies. As expected, CodY was detected only in the protoplast and WCL fractions, and was completely absent in the CW samples, confirming that this fraction was free of cytoplasmic contaminants **(Figure 3).** Importantly, the PepNFLAG protein was clearly detected in the both the WCL and CW fractions, with a fainter band present in the protoplast sample **(Figure 3)**. Thus, these data suggest that despite the absence of a canonical LPxTG motif, Sec-dependent signal peptide, as well as any known cell wall binding motif, PepN indeed localizes to the CW compartment.

**Figure 3.**
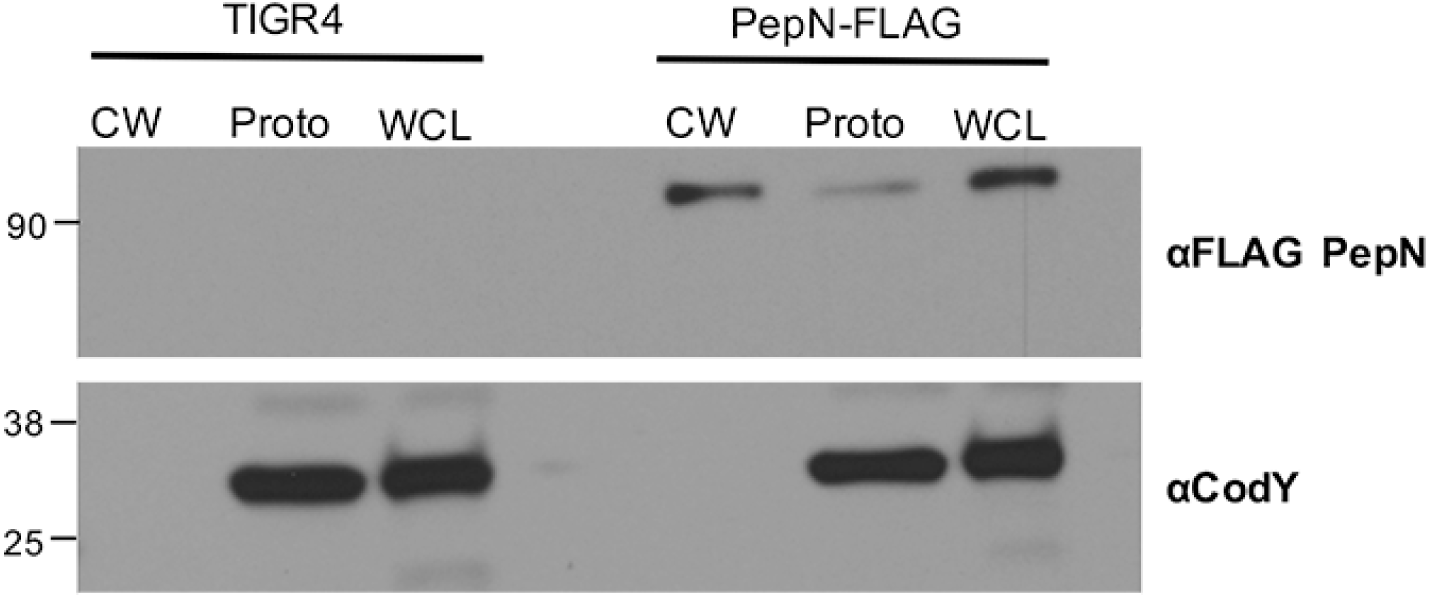
PepN localizes to the cell wall compartment within *Spn* cells. TIGR4 and PepNFLAG strains were grown to mid-log phase and cells were collected via centrifugation. Cells were treated with CWDB for 2h at 37°C with rotation, followed by centrifugation using a 0.22μm spin column to separate the CW and protoplast fractions. Whole cell lysates (WCL) were prepared by adding sample buffer directly to whole cells followed by a 10 min incubation at 100°C. Samples were then analyzed by Western blotting using anti-FLAG or anti-CodY (a cytoplasmic protein) antibodies. Data shown is from one experiment representative of three independent experiments.

### Neutrophil elastase degrades PepN in intact *Spn* cells

To validate PepN as NE substrate and confirm that NE can degrade PepN in the context of native cell wall architecture, we exposed ~5 × 10^6^ wild-type or PepNFLAG cells to 6.8μM NE, followed by subcellular fractionation and Western Blot analysis using anti-FLAG or anti-CodY antibodies. Again, the CodY signal was only present in the WCL samples and the intensity of this band was unchanged in the presence of NE. Strikingly, analysis of the CW and WCL fractions revealed a strong PepNFLAG signal in the untreated samples that was markedly diminished upon exposure to NE **(Figures 4A and B)**. These data strongly suggest that in the context of an intact *Spn* cell, PepN is a *bona fide* CW protein that is markedly degraded by NE.

**Figure 4.**
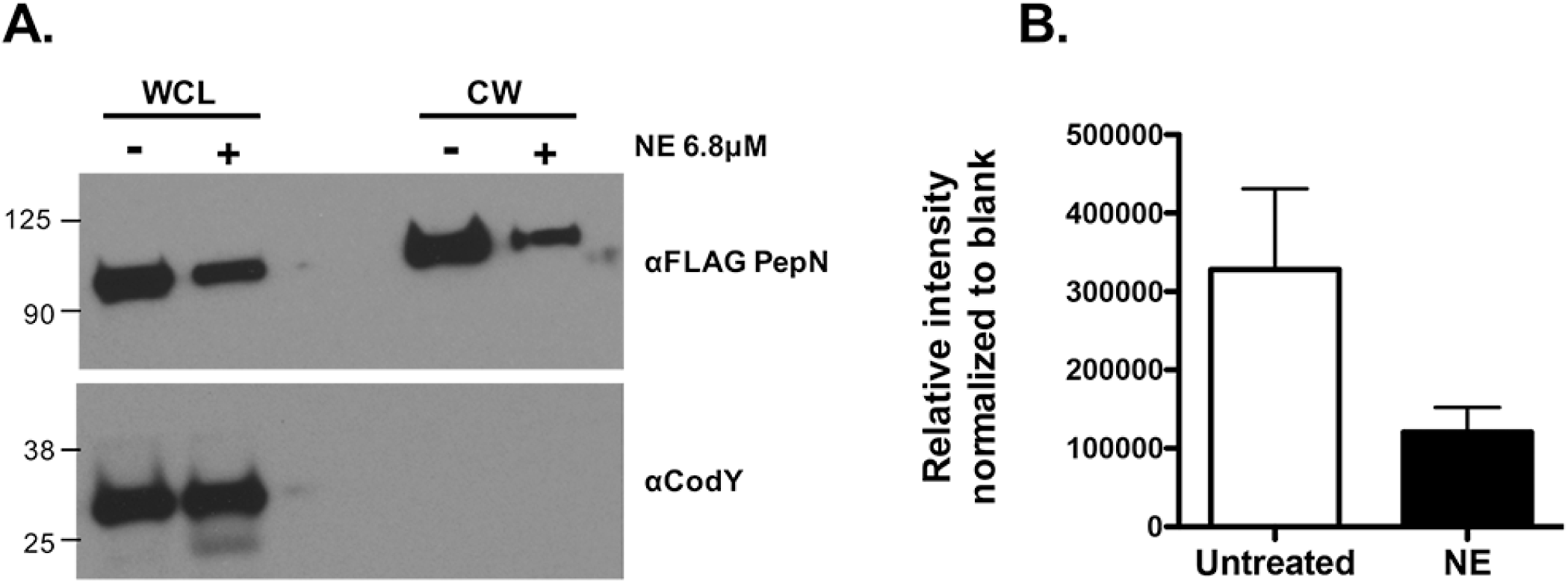
NE degrades PepN in intact *Spn* cells. TIGR4 and PepNFLAG strains were grown to early-log phase in TSB supplemented with catalase and cells were either left untreated or were exposed to 6.8μM NE for 1h at 37°C. **(A)** Cells were fractionated to isolate the CW and samples were analyzed via Western blotting using anti-FLAG or anti-CodY (a cytoplasmic protein) antibodies. Data shown is from one experiment representative of four independent experiments. (**B)** Band intensity in CW samples were quantified using ImageStudioLite software. Compared to the untreated control, PepN intensity was reduced ~3-fold in NE-treated samples. Data presented are the means ± SD from 4 independent experiments.

### The Δ*pepN* mutant demonstrates enhanced resistance to killing by purified NE *in vitro* and by human neutrophils *ex vivo*

To assess the contribution of PepN to the effective killing of *Spn in vitro,* we generated a mutant strain lacking this protein (*ΔpepN*). To first characterize this mutant, we assessed its growth kinetic phenotype, and determined that it was similar to that of wild-type TIGR4 (**Figure 5A)**. Additionally, to ensure that removal of the cell wall protein PepN did not have an unexpected impact on capsule level, a structure that was demonstrated to influence susceptibility to NSPs [37], we quantified and compared capsule on wild-type and Δ*pepN* cells. As shown in **Figures 5B and 5C,** while wild-type TIGR4 and *ΔpepN* cells express significantly more capsule than the acapsular control (Δ*cps*), there was no significant difference between wild-type TIGR4 and the *ΔpepN* strain.

**Figure 5.**
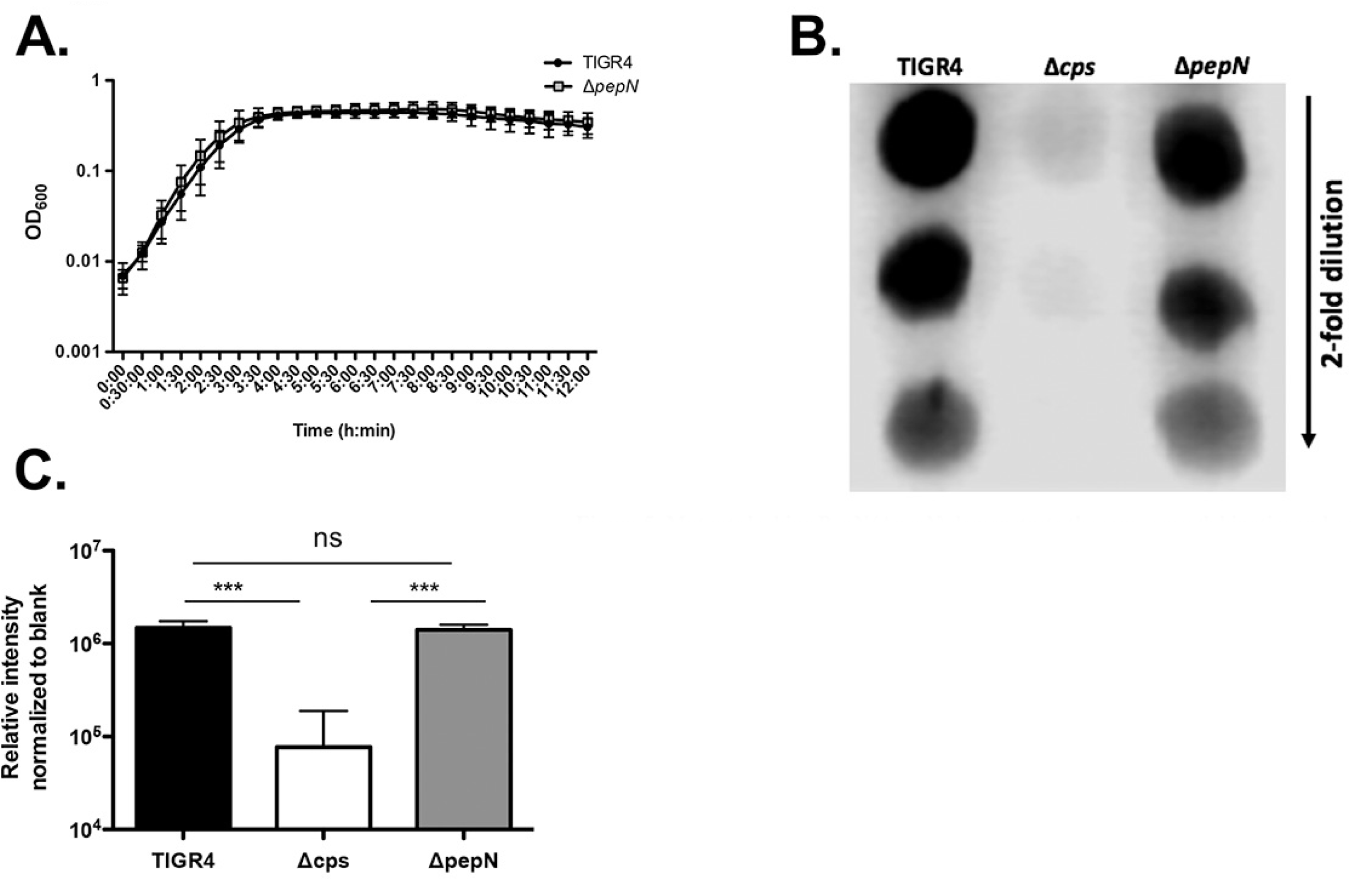
Mutants lacking PepN (Δ*pepN)* demonstrate the same growth kinetics and capsule levels as wild-type TIGR4. A) Mutants lacking PepN (Δ*pepN)* demonstrate the same growth kinetics and capsule expression levels as wild-type TIGR4. A) TIGR4 and Δ*pepN* strains of *S. pneumoniae* were grown to mid-log phase in THY supplemented with oxyrase. Cells were then back-diluted in fresh THY/oxyrase to OD_600_ 0.003 and 200μL of cells were plated in replicates of 8 wells into 96 well plates. Cells were incubated at 37°C and OD_600_ was measured every 30 minutes for 12h. Data presented are the means ± SD from 3 independent experiments. **B and C)** TIGR4, Δ*cps* and *ΔpepN* cells were grown to mid-log phase in THY supplemented with oxyrase. 1mL of cells were isolated, washed in 1X PBS and resuspended in an appropriate volume of 1X PBS to match OD across all three strains. A 300μL aliquot of each strain was pelleted, resuspended in 300μL CWDB and incubated for 30 mins at 37°C. Samples were then sonicated for 4 – 10 sec intervals while on ice and subsequently serially diluted in 1X PBS. For each strain, 5μL spots were applied to a nitrocellulose membrane with suction, followed by probing with an unconjugated rabbit anti-serotype 4 serum and HRP-conjugated goat anti-rabbit antibody. Membranes were then developed using ECL Blotting Substrate and visualized using the C-Digit Western Blot Scanner. **B)** Data shown are from one blot representative of 3 independent experiments. **C)** Data presented are the means ± SD from 3 independent experiments. P<0.001 for both comparisons between TIGR4 and Δ*cps* and Δ*pepN* and Δ*cps*. There was no significant difference between TIGR4 and Δ*pepN*. Statistical analyses were conducted using a one-way ANOVA and Tukey’s Multiple Comparisons Test.

In subsequent experiments, we assessed the phenotype of the *ΔpepN* strain in the presence of various concentrations of purified NE. These data demonstrated that the *ΔpepN* mutant was significantly more resistant to NE-mediated killing as compared to wild-type TIGR4 (**Figures 6A and B)**. Importantly, as shown in **Figure 6B,** when we reverted the *ΔpepN* mutant (*ΔpepN*Rev) and exposed this strain to 3.4μM purified NE, it exhibited a sensitive phenotype similar to that of wild-type TIGR4. To test the *ΔpepN* mutant in a more physiologically relevant setting, we determined its survival phenotype using an opsonophagocytic assay with purified human neutrophils. These experiments revealed that, compared to wild-type *Spn,* the *ΔpepN* mutant was also significantly more resistant to phagocytic killing by human neutrophils **(Figure 6C)**. In fact, while wild-type TIGR4 was killed in the presence of PMNs, the *ΔpepN* mutant was not and displayed very slight growth **(Figure 6C)**. Taken together, these data reveal that the nonessential PepN protein is not only a substrate of NE, but also that its degradation plays an important role in the effective killing of *Spn* by both purified NE and whole human neutrophils.

**Figure 6.**
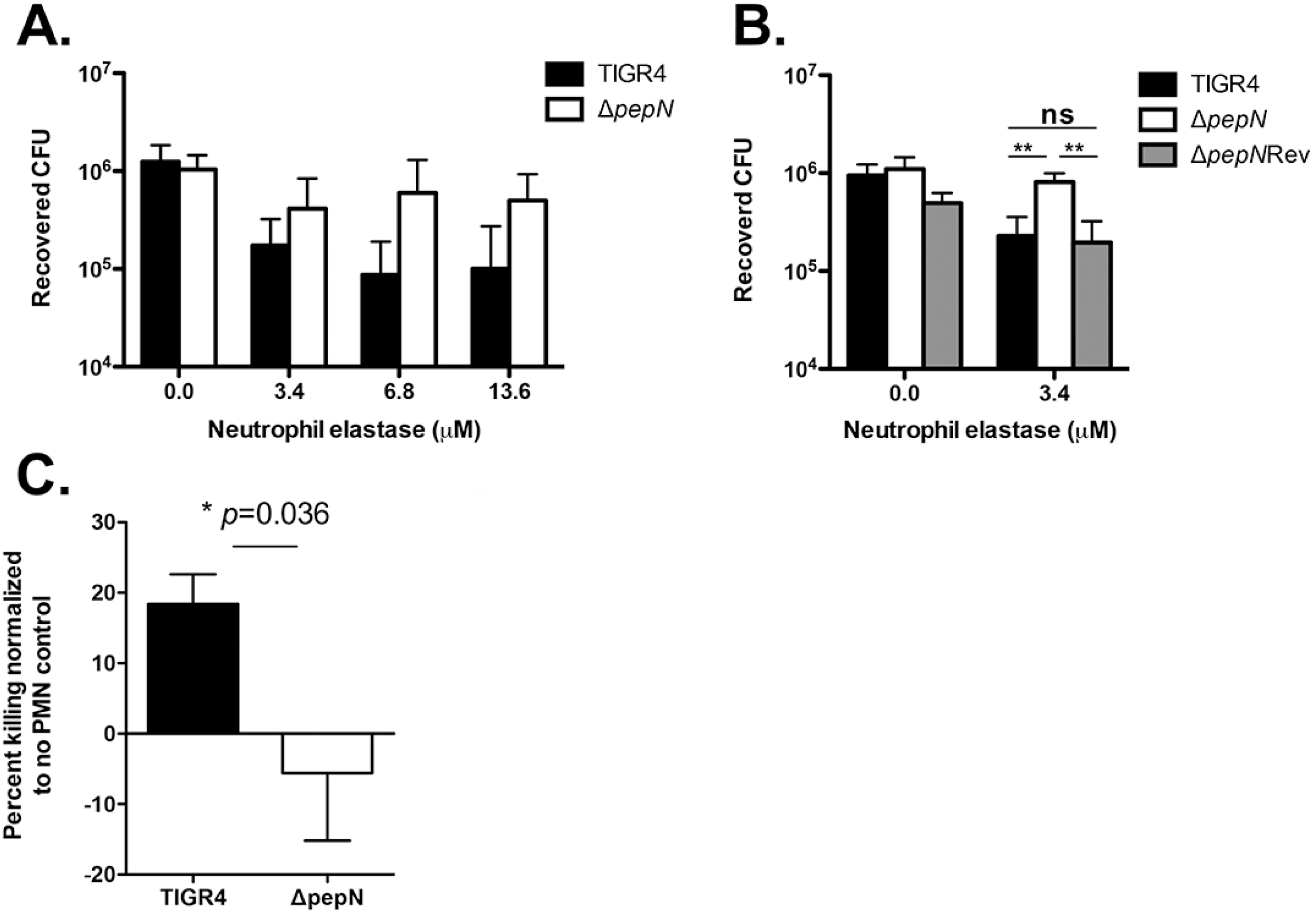
The Δ*pepN* mutant demonstrates enhanced resistance to NE-mediated killing *in vitro* and to killing by human neutrophils *ex vivo*. For *in vitro* bactericidal assays using NE, strains were grown to early-log phase in TSB supplemented with catalase, cells were collected by centrifugation, resuspended in 50mM sodium phosphate and ~5 × 10^5^ CFU were treated with various concentrations of NE or left untreated for 1h at 37°C. Samples were then serially diluted and plated to **A)** enumerate viable CFU. P=0.004 comparing TIGR4 to *ΔpepN* across all concentrations of NE and was calculated using two-way ANOVA. Data presented are the means ± SD from 3-4 independent experiments. **B)** TIGR4, Δ*pepN* and *ΔpepNRev* cells were exposed to 3.4μM NE or left untreated for 1h at 37°C, followed by serial dilution and enumeration of CFU to calculate percent survival relative to the paired untreated control. P<0.01 comparing TIGR4 to Δ*pepN* and Δ*pepN* to *ΔpepN*Rev; P=ns comparing TIGR4 to *ΔpepN*Rev. P values were calculated using one-way ANOVA. Data presented are the means ± SD from 3 independent experiments. **C)** For *ex vivo* opsonophagocytic killing experiments, PMNs were isolated from the blood of healthy donors and incubated with pre-opsonized *Spn* for 45 mins at 37°C with rotation. Reactions were then stopped by placing samples on ice and viable CFU were determined after serial dilution and plating. The percentage of bacteria killed upon incubation with PMNs was determined by comparing surviving CFU for each strain to a no PMN control. Positive percent killing indicates bacterial death while negative percent indicates bacterial growth. P=0.036 and was calculated using student’s t-test. Data are shown as the means ± SD pooled from 4 independent experiments with PMNs from four separate donors.

## DISCUSSION

In neutrophils, non-oxidative mechanisms of killing phagocytosed *Spn* were shown to be essential both *in vitro* and *in vivo* and primarily involve the activity of NSPs, such as NE and CG [9, 10]. Further supporting the importance of NSPs, individuals with impaired granule mobilization, which affects the fusion of NSP-containing granules with the phagolysosome, exhibit a defect in neutrophil-mediated killing of *Spn* [12]. In addition to playing a key role in controlling *Spn* infections, other studies demonstrated that NSPs have the capacity to directly kill or destroy virulence factors produced by a variety of pathogens including *P. aeruginosa, E. coli, S. flexneri, K. pneumoniae* and *S. aureus* [18-23]. Although previous reports highlight the importance of NSPs in bacterial clearance, little is known about the identity of the *Spn* proteins that are degraded by NSPs and facilitate effective killing.

In this study, we aimed to identify *Spn* CW proteins that are degraded by the NSPs NE and CG. To do so, we exposed a purified CW preparation to NE and CG *in vitro* and observed the degradation of a ~90kDa protein only by NE, suggesting a difference in substrate specificity between these two NSP family members. Subsequent analysis via mass spectrometry revealed this ~90kDa protein to be aminopeptidase N (PepN). Furthermore, these experiments demonstrated that the relative abundance of PepN was reduced 10^3^-10^4^-fold in purified CW samples treated with NE as compared to untreated controls, indicating that NE specifically and substantially degrades this *Spn* protein. Interestingly, in the context of an intact *Spn* cell, PepN abundance was more modestly reduced (~3-fold) in NE-treated samples, as compared to untreated controls. This discrepancy may be due, in part, to the masking effect provided by the capsular polysaccharide and the complex architecture of the cell wall, thus limiting the accessibility of PepN as a NE substrate. Nevertheless, degradation of PepN on intact *Spn* cells was still possible and sufficient to induce bacterial cell death.

Since analysis of the PepN sequence failed to reveal a canonical Sec-dependent secretion signal, LPxTG cell wall anchoring motif or other export signals [35, 36], we created a mutant strain of *Spn* harboring a FLAG-tagged version of the protein to more definitively assess its subcellular localization. Interestingly, despite the absence of an obvious secretory or CW localization motif, the majority of PepN was identified in the CW compartment. Although these findings are at odds with the reported cytoplasmic localization of PepN in *S. mitis* (97% sequence identity) and *S. salivarius* (67% sequence identity) [38, 39], the absence of the cytoplasmic control protein, CodY, in our CW fractions provides confidence that PepN is indeed cell wall localized in *Spn*. Additionally, in various other Gram-positive species including *L. lactis,* PepN (57% identity) is a CW anchored protein, indicating there is some variability in the localization of this protein [40]. Thus, akin to other *Spn* proteins including pneumolysin [30], PavA [41] and HtrA [35], our data indicate that PepN is yet another example of a non-classical protein that localizes to the cell wall despite the absence of an obvious export sequence.

These initial experiments strongly suggest that PepN is a CW-localized protein that serves as a substrate for NE. However, these conclusions were drawn from experiments using a purified preparation of CW proteins, which may affect the abundance, diversity or availability of substrates for NE-mediated degradation. More importantly, since the aim of this study was to identify *Spn* proteins that not only were degraded by NSPs, but were also involved in NSP-mediated killing of *Spn,* we conducted additional experiments in a more physiologically relevant system. Through experiments that exposed intact *Spn* cells expressing the FLAG-tagged version of PepN to NE, we confirmed that PepN does indeed localize to the CW and it is markedly degraded compared to untreated controls. Previous studies in *E. coli* and *P. aeruginosa* identified the outer membrane proteins OmpA and OprF, respectively, as targets that are degraded by NE that results in bacterial cell death [20, 22]. Both of these proteins are porins that contribute to virulence and isogenic mutants lacking either OmpA or OprF in the respective strain demonstrated enhanced resistance to NE-mediated killing *in vitro* and in relevant mouse models of infection [20, 22, 42, 43]. To directly assess the contribution of PepN to NSP-mediated killing of *Spn* cells, we created the Δ*pepN* strain and confirmed that its growth rate was indistinguishable from that of wild-type. Importantly, since capsule level was shown to impact killing by NSPs [37], and it was feasible that the deletion of a CW-localized aminopeptidase could affect the attachment of capsule to the cell, we quantified capsule on wild-type TIGR4 and Δ*pepN* cells. Data from these experiments and India Ink staining (data not shown), demonstrated that both strains possess similar amounts of capsule. *In vitro* bactericidal assays that exposed wild-type TIGR4 and Δ*pepN* cells to purified NE revealed that cells lacking PepN were significantly more resistant to killing. Additionally, Δ*pepN* cells were significantly more resistant to opsonophagocytic killing by whole human neutrophils *ex vivo*. Based on the observation that Δ*pepN* cells express wild-type levels of capsule, we speculate that the differential killing phenotype is not attributed to variations in C3 deposition [44-46], but rather due to the absence of a *bona fide* NE substrate that is necessary for optimal killing of *Spn* once in the phagolysosome.

The key observation from these experiments was that NE-mediated degradation of a non-essential CW protein, PepN, was sufficient to induce *Spn* killing. One possible explanation for this observation is that degradation of PepN is sufficient to destabilize the cell envelope and induce cell lysis. Alternatively, if PepN is attached, in some fashion, to peptidoglycan, teichoic or lipotechoic acid, its degradation by NE may also damage these essential structures or impair their turnover such that normal *Spn* growth is disrupted. Another potential explanation may be that, via its aminopeptidase activity, PepN modifies other CW proteins and creates additional NE substrates. Thus in PepN-sufficient *Spn* cells, not only is PepN directly degraded, but also the modified CW proteins created via PepN aminopeptidase activity may also be degraded, which together is enough to cause cell death. It would be possible to test this last hypothesis by creating a mutant strain with an inactivated PepN catalytic site and assess its viability in the presence of NE. These unanswered questions are beyond the scope of this current study, but emphasize the need for future experiments.

The role of PepN in *Spn* biology and in the context of host infection is not well understood. However, in other closely related species of *Streptococcus*, including *S. thermophilus,* PepN is characterized as a 95kDa monomeric, metallo-aminopeptidase that possesses three Zn^2+^ binding sites that coordinates its active site [24, 40]. Based on its sequence homology to other aminopeptidases, it is thought that PepN functions to degrade endogenous proteins and contributes to normal protein turnover as well as to provide free amino acids to be used for metabolic processes [40]. Interestingly, a recent study demonstrated that PepN present in *Spn* lysates dampens the effector function of cytotoxic T lymphocyte by modulating the intracellular TCR signaling cascade and ultimately dampens the production of the pro-inflammatory cytokine, IFN-γ [47]. Together with the findings in our study, these data begin to reveal a previously underappreciated role for PepN in host-pathogen interactions and *Spn* disease pathogenesis. In summary, this is the first report to identify a *Spn* protein, PepN, as a substrate that is degraded by NE and that also plays a key role in NSP-mediated killing of *Spn* both *in vitro* and *ex vivo.* Additionally, we determined that despite the absence of a canonical export sequence, PepN localizes to the CW compartment within *Spn.* We propose a model where *Spn* cells are opsonophagocytosed by neutrophils and are subsequently bombarded with an assortment of lysosome- and granule-derived anti-microbial factors. Based on our data, a key step in this process is the NE-mediated degradation of PepN, which contributes to the effective killing of *Spn*.

## ACKNOWLEDGEMENTS

This project was funded, in part, by Undergraduate Research Awards provided by the Jess and Mildred Fisher College of Science and Mathematics at Towson University. We thank Dr. A. L. Sonenshein for providing the anti-CodY antibody and Dr. John Leong for sharing additional reagents needed for PMN experiments. We also thank Neil Greene for helpful discussions and for critically reading our manuscript.

## REFERENCES

1. Kadioglu, A., et al., The role of Streptococcus pneumoniae virulence factors in host respiratory colonization and disease. Nat Rev Microbiol, 2008. 6(4): p. 288–301.

2. Mitchell, T.J., Virulence factors and the pathogenesis of disease caused by Streptococcus pneumoniae. Res Microbiol, 2000. 151(6): p. 413–9.

3. Tuomanen, E.I., R. Austrian, and H.R. Masure, Pathogenesis of pneumococcal infection. N Engl J Med, 1995. 332(1): p. 1280–4.

4. Weiser, J.N., The pneumococcus: why a commensal misbehaves. J Mol Med, 2009. 88(2): p. 97–102.

5. Pneumococcal Disease. 2017; Available from: http://www.who.int/ith/diseases/pneumococcal/en/.

6. Dallaire, F., et al., Microbiological and inflammatory factors associated with the development of pneumococcal pneumonia. J Infect Dis, 2001. 184(3): p. 292–300.

7. van Rossum, A.M., E.S. Lysenko, and J.N. Weiser, Host and bacterial factors contributing to the clearance of colonization by Streptococcus pneumoniae in a murine model. Infect Immun, 2005. 73(1): p. 7718–26.

8. Segal, A.W., How neutrophils kill microbes. Annu Rev Immunol, 2005. 23: p. 197–223.

9. Standish, A.J., Weiser, J.N., Human Neutrophils Kill Streptococcus pneumoniae via Serine Proteases. Journal of Immunology, 2009. 183(4): p. 2602–2609.

10. Hahn, I., et al., Cathepsin G and neutrophil elastase play critical and nonredundant roles in lung-protective immunity against Streptococcus pneumoniae in mice. Infect Immun, 2011. 79(1): p. 4893–901.

11. Ganz, T., et al., Microbicidal/cytotoxic proteins of neutrophils are deficient in two disorders: Chediak-Higashi syndrome and “specific” granule deficiency. J Clin Invest, 1988. 82(2): p. 552–6.

12. Root, R.K., A.S. Rosenthal, and D.J. Balestra, Abnormal bactericidal, metabolic, and lysosomal functions of Chediak-Higashi Syndrome leukocytes. J Clin Invest, 1972. 51(3): p. 649–65.

13. Perera, N.C., et al., NSP4, an elastase-related protease in human neutrophils with arginine specificity. Proc Natl Acad Sci U S A, 2012. 109(1): p. 6229–34.

14. Korkmaz, B., T. Moreau, and F. Gauthier, Neutrophil elastase, proteinase 3 and cathepsin G: physicochemical properties, activity and physiopathological functions. Biochimie, 2008. 90(2): p. 227–42.

15. Hellman, L. and M. Thorpe, Granule proteases of hematopoietic cells, a family of versatile inflammatory mediators - an update on their cleavage specificity, in vivo substrates, and evolution. Biol Chem, 2014. 395(1): p. 15–49.

16. Borregaard, N. and J.B. Cowland, Granules of the human neutrophilic polymorphonuclear leukocyte. Blood, 1997. 89(1): p. 3503–21.

17. Korkmaz, B., et al., Neutrophil elastase, proteinase 3, and cathepsin G as therapeutic targets in human diseases. Pharmacol Rev, 2010. 62(4): p. 726–59.

18. Weinrauch, Y., et al., Neutrophil elastase targets virulence factors of enterobacteria. Nature, 2002. 417(6): p. 91–4.

19. Hazenbos, W.L., et al., Novel staphylococcal glycosyltransferases SdgA and SdgB mediate immunogenicity and protection of virulence-associated cell wall proteins. PLoS Pathog, 2013. 9(1): p. e1003653.

20. Belaaouaj, A., K.S. Kim, and S.D. Shapiro, Degradation of outer membrane protein A in Escherichia coli killing by neutrophil elastase. Science, 2000. 289(5): p. 1185–8.

21. Belaaouaj, A., et al., Mice lacking neutrophil elastase reveal impaired host defense against gram negative bacterial sepsis. Nat Med, 1998. 4(5): p. 615–8.

22. Hirche, T.O., et al., Neutrophil elastase mediates innate host protection against Pseudomonas aeruginosa. J Immunol, 2008. 181(7): p. 4945–54.

23. Lopez-Boado, Y.S., et al., Neutrophil serine proteinases cleave bacterial flagellin, abrogating its host response-inducing activity. J Immunol, 2004. 172(1): p. 509–15.

24. Chavagnat, F., M.G. Casey, and J. Meyer, Purification, characterization, gene cloning, sequencing, and overexpression of aminopeptidase N from Streptococcus thermophilus A. Appl Environ Microbiol, 1999. 65(7): p. 3001–7.

25. Heckman, K.L. and L.R. Pease, Gene splicing and mutagenesis by PCR-driven overlap extension. Nat Protoc, 2007. 2(4): p. 924–32.

26. Hava, D.L. and A. Camilli, Large-scale identification of serotype 4 Streptococcus pneumoniae virulence factors. Mol Microbiol, 2002. 45(5): p. 1389–406.

27. Salles, C., et al., The high level streptomycin resistance gene from Streptococcus pneumoniae is a homologue of the ribosomal protein S12 gene from Escherichia coli. Nucleic Acids Res, 1992. 20(2): p. 6103.

28. Dalia, A.B., E. McDonough, and A. Camilli, Multiplex genome editing by natural transformation. Proc Natl Acad Sci U S A, 2014. 111(2): p. 8937–42.

29. Bricker, A.L. and A. Camilli, Transformation of a type 4 encapsulated strain of Streptococcus pneumoniae. FEMS Microbiol Lett, 1999. 172(2): p. 131–5.

30. Price, K.E. and A. Camilli, Pneumolysin localizes to the cell wall of Streptococcus pneumoniae. J Bacteriol, 2009. 191(7): p. 2163–8.

31. Hurley, B.P., et al., Polymorphonuclear cell transmigration induced by Pseudomonas aeruginosa requires the eicosanoid hepoxilin A3. J Immunol, 2004. 173(9): p. 5712–20.

32. Bou Ghanem, E.N., et al., The Alpha-Tocopherol Form of Vitamin E Boosts Elastase Activity of Human PMNs and Their Ability to Kill Streptococcus pneumoniae. Front Cell Infect Microbiol, 2017. 7: p. 161.

33. Dalia, A.B., A.J. Standish, and J.N. Weiser, Three surface exoglycosidases from Streptococcus pneumoniae, NanA, BgaA, and StrH, promote resistance to opsonophagocytic killing by human neutrophils. Infect Immun, 2010. 78(5): p. 2108–16.

34. Zar, J.H., Biostatistical Analysis. 5^th^ ed. 2010: Pearson.

35. Perez-Dorado, I., S. Galan-Bartual, and J.A. Hermoso, Pneumococcal surface proteins: when the whole is greater than the sum of its parts. Mol Oral Microbiol, 2012. 27(4): p. 221–45.

36. Plumptre, C.D., A.D. Ogunniyi, and J.C. Paton, Surface association of Pht proteins of Streptococcus pneumoniae. Infect Immun, 2013. 81(1): p. 3644–51.

37. van der Windt, D., et al., Nonencapsulated Streptococcus pneumoniae resists extracellular human neutrophil elastase- and cathepsin G-mediated killing. FEMS Immunol Med Microbiol, 2012. 66(3): p. 445–8.

38. Andersson, C., et al., Purification and characterization of an aminopeptidase from Streptococcus mitis ATCC 903. Curr Microbiol, 1992. 25(5): p. 261–7.

39. Midwinter, R.G. and G.G. Pritchard, Aminopeptidase N from Streptococcus salivarius subsp. thermophilus NCDO 573: purification and properties. J Appl Bacteriol, 1994. 77(3): p. 288–95.

40. Gonzales, T. and J. Robert-Baudouy, Bacterial aminopeptidases: properties and functions. FEMS Microbiol Rev, 1996. 18(4): p. 319–44.

41. Holmes, A.R., et al., The pavA gene of Streptococcus pneumoniae encodes a fibronectinbinding protein that is essential for virulence. Mol Microbiol, 2001. 41(6): p. 1395–408.

42. Weiser, J.N. and E.C. Gotschlich, Outer membrane protein A (OmpA) contributes to serum resistance and pathogenicity of Escherichia coli K-1. Infect Immun, 1991. 59(7): p. 2252–8.

43. Woodruff, W.A., et al., Expression in Escherichia coli and function of Pseudomonas aeruginosa outer membrane porin protein F. J Bacteriol, 1986. 167(2): p. 473–9.

44. Hyams, C., et al., The Streptococcus pneumoniae capsule inhibits complement activity and neutrophil phagocytosis by multiple mechanisms. Infect Immun, 2010. 78(2): p. 704–15.

45. Hyams, C., et al., Streptococcus pneumoniae resistance to complement-mediated immunity is dependent on the capsular serotype. Infect Immun, 2009. 78(2): p. 716–25.

46. Shainheit, M.G., et al., Mutations in pneumococcal cpsE generated via in vitro serial passaging reveal a potential mechanism of reduced encapsulation utilized by a conjunctival isolate. J Bacteriol, 2015. 197(1): p. 1781–91.

47. Blevins, L.K., et al., A Novel Function for the Streptococcus pneumoniae Aminopeptidase N: Inhibition of T Cell Effector Function through Regulation of TCR Signaling. Front Immunol, 2017. 8: p. 1610.

